# Insulin-independent glucose uptake in skeletal muscle by coupled SGLT and Na,K-ATPase transport

**DOI:** 10.64898/2026.03.24.714065

**Authors:** Natalie J. Norman, Tatiana L. Radzyukevich, Peter W. Chomczynski, Michal Rymaszewski, Izabela Fokt, Waldemar Priebe, Lucas Schmidt, Tiffany Zhu, Bryan Mackenzie, Julio A. Landero, Judith A. Heiny

## Abstract

Exercise is a cornerstone therapy for diabetes because working skeletal muscles take up glucose at dramatically greater rates than postprandial insulin-stimulated glucose uptake and, notably, do so without a requirement for insulin. This remarkable ability of working muscles is preserved in diabetes, when muscles become resistant to insulin. However, the mechanism of insulin-*in*dependent glucose uptake by working muscles is not fully understood. Here we describe a previously unrecognized glucose uptake pathway in muscle, which we refer to as “mSGLT” based on shared properties with the Sodium Glucose Linked Transporter family. In contrast to the abundant GLUT4 transporter, mSGLT is not regulated by insulin, requires Na,K-ATPase-α2 activity, and transports the hexose α-methyl-D-glucoside (αMDG), a glucose derivative that is handled by SGLTs but not GLUT4. The mSGLT pathway and GLUT transport pathways are independent and additive. In addition to exercise, mSGLT imports glucose under other conditions of adrenergic stimulation, which inhibits pancreatic insulin release and reduces the insulin sensitivity of muscle. SGLT2-specific antibodies recognize a protein in muscle of similar size to the kidney SGLT2; this protein localizes to the muscle t-tubules, together with Na,K-ATPase-α2 and MAP17, the regulatory subunit of SGLT2. However, skeletal muscles do not express a full-length transcript of S*lc5a2* (SGLT2), and SGLT2-specific inhibitors do not inhibit mSGLT with high affinity. The novel transporter may be a muscle variant of *Slc5a2* that results from post-transcriptional or post-translational mechanisms. mSGLT and its regulation offer potential muscle-specific therapeutic targets for treating hyperglycemia and other conditions when insulin-stimulated glucose disposal into muscle is impaired.

## INTRODUCTION

The skeletal musculature is the major depot for glucose storage and utilization and a primary regulator of glucose homeostasis ^1^. Muscles use both insulin-mediated and non-insulin-mediated pathways to import glucose from blood and need both to achieve glucose homeostasis. Insulin-stimulated glucose uptake predominates after a meal when circulating insulin rises and stimulates muscle to take up and store approximately 80% of the ingested glucose. Postprandial insulin stimulation approximately doubles the muscle capacity for glucose uptake by increasing the membrane density of GLUT4 (*Slc2a4*) ^1^, the insulin-regulated glucose transporter that is highly expressed in skeletal muscle. This mechanism uses a well-described pathway whereby insulin binding to muscle insulin receptors induces translocation of GLUT4 from intracellular stores to the plasma membrane, mainly the t-tubules. ^1-6^ The imported glucose is converted to glycogen and stored until needed for muscle contraction

Non-insulin mediated glucose uptake occurs during periods of declining insulin and elevated catecholamines. These conditions occur acutely during exercise and chronically during periods of stress, after injury, and with overnutrition and obesity. Epinephrine and norepinephrine released in these conditions inhibit insulin secretion from pancreatic beta cells and reduce the insulin sensitivity of muscle and fat. ^2^ Non-insulin mediated glucose uptake rates by muscle are significant; during intense exercise, they can exceed postprandial uptake by 50-100-fold. ^3,4^ Non-insulin-mediated glucose uptake is also important in metabolic disorders such as insulin resistance and diabetes, when insulin-stimulated uptake is impaired. In these conditions, non-insulin-mediated glucose uptake contributes up to 70% of total glucose disposal. ^5^

Despite its importance and large capacity, the molecular mechanisms that underlie non-insulin-mediated glucose transport into muscle are not fully understood. Muscles can import glucose without insulin, in part, using a non-insulin triggered mode of GLUT4 translocation. Contraction-related factors, not yet fully identified, are able to trigger GLUT4 translocation using a signaling pathway that bypasses the muscle insulin receptor but converges at intracellular nodes shared with the canonical insulin signaling network.^1^ The actions of insulin and contraction on GLUT4 translocation are additive ^6,7^, On the other hand, studies of non-insulin triggered GLUT4 translocation have not fully explained puzzling inconsistencies found with GLUT4 knockout models. The absence of an essential glucose transporter that “does it all” was expected to produce a severe metabolic phenotype. However, a global GLUT4 knockout model did not develop diabetes ^8^ and mice with a muscle specific deletion of GLUT4 show either normal insulin tolerance and glucose uptake or impaired glucose uptake with insulin resistance, depending on model and genetic background ^9-13^. These variable phenotypes have led to the proposal that additional mechanisms of non-insulin mediated glucose uptake may exist and contribute differently to the different models. The finding that non-insulin mediated glucose uptake remains when GLUTs 1, 3, 4, 6 and 10 are inhibited or GLUT1 is depleted seems to exclude known GLUT subtypes as the candidate transporter ^14^.

This study tests the hypothesis that a sodium-glucose-linked transporter (SGLT) may also contribute to non-insulin mediated glucose uptake in muscle. SGLTs, the other major family of glucose transporters, use the inward Na gradient generated by Na,K-ATPase activity to drive glucose together with Na^+^ into cells ^15^. Muscles use Na-linked transport to import many essential ions and nutrients including Na-PO^4^ uptake, Na-Ca exchange, and Na-amino acid uptake ^16^. Muscle contraction dramatically stimulates Na,K-ATPase-α2, the predominant pump isoform in skeletal muscle, during and immediately following contraction ^17-22^. The resulting, steepened Na gradient drives nutrient uptake to supply the needs of contraction and replenish energy stores.

To test this hypothesis, we measured simultaneous glucose transport and Na,K-ATPase activity in isolated, mouse EDL muscles at rest, during and post contraction, and during adrenergic stimulation. We measured uptake of sugars and Rb, a congener for K^+^ transport by Na,K-ATPase, using Inductively Coupled Plasma-Mass Spectroscopy (ICP-MS) ^23-25^. We used naturally abundant RbCl at tracer concentrations to measure Na,K-ATPase transport and ^13^C carbon-substituted hexoses to measure glucose uptake. We identified GLUT4 and SGLT uptake using glucose derivatives that are specific substrates for GLUT4 or SGLTs. Both GLUT4 and SGLTs transport (6-^13^C-glucose, hereafter glucose), the natural substrate of all glucose transporters. GLUT4 selectively transports 6-^13^C-2-deoxy-D-glucose (hereafter 2-DG) and SGLTs selectively transport 6-^13^C αMDG (6-^13^C-αMDG, hereafter αMDG). ^15^

## RESULTS

### A component of glucose uptake in contracting muscle requires Na,K-ATPase-α2 activity

To investigate whether a Na-glucose linked transporter contributes to contraction-stimulated glucose uptake, we measured the simultaneous uptake of Rb and ^13^C-6-carbon-substituted hexoses in isolated EDL muscles at rest and during contraction under nominally insulin-free conditions (Fig. 1a-1c). We compared uptake of glucose, which reports total glucose uptake by all glucose transporters, with uptake of αMDG, a glucose derivative transported by SGLTs but not GLUT4. Na,K-ATPase activity generates the inward Na gradient used by SGLTs to import glucose together with Na^+^ into cells. Rb is a congener for K transport by Na,K-ATPase. Because αMDG is not metabolized by glycolysis, this measurement captures all ^13^C that enters the muscle during the uptake period without loss to CO_2_. The EDL muscle uses carbohydrate as the predominant substrate for generating ATP; without a metabolizable substrate, glucose-6-phosphate from glycogenolysis is the sole substrate for glycolysis in this protocol. Nominally insulin-free conditions were achieved by lowering serum insulin with fasting, using isolated muscles removed from the circulation, perfusing the muscle for 30 min in insulin-free buffer, and omitting insulin from all solutions. The stimulation pattern was designed to strongly activate Na,K-ATPase-α2. Na,K-ATPase-α2 transport was identified as the component of Rb uptake that is inhibited by 0.75 μM ouabain. In rodents, the Na,K-ATPase-α2 pump is the sole pump isoform sensitive to inhibition by μM ouabain ^26^. This concentration of ouabain does not significantly alter the resting membrane potential of muscle ^27^, and it did not significantly change muscle force or development of fatigue (Fig. 1c). Therefore, differences in sugar uptake between control and ouabain-treated muscles cannot be attributed to differences in contractility. Membrane potential exerts a small effect on the transport rate of some SGLTs. A negative membrane potential favors greater transport rates of vSGLT ^28^. In this stimulation protocol, the membrane potential is negative for ∼95% of the uptake period.

**Fig. 1.**
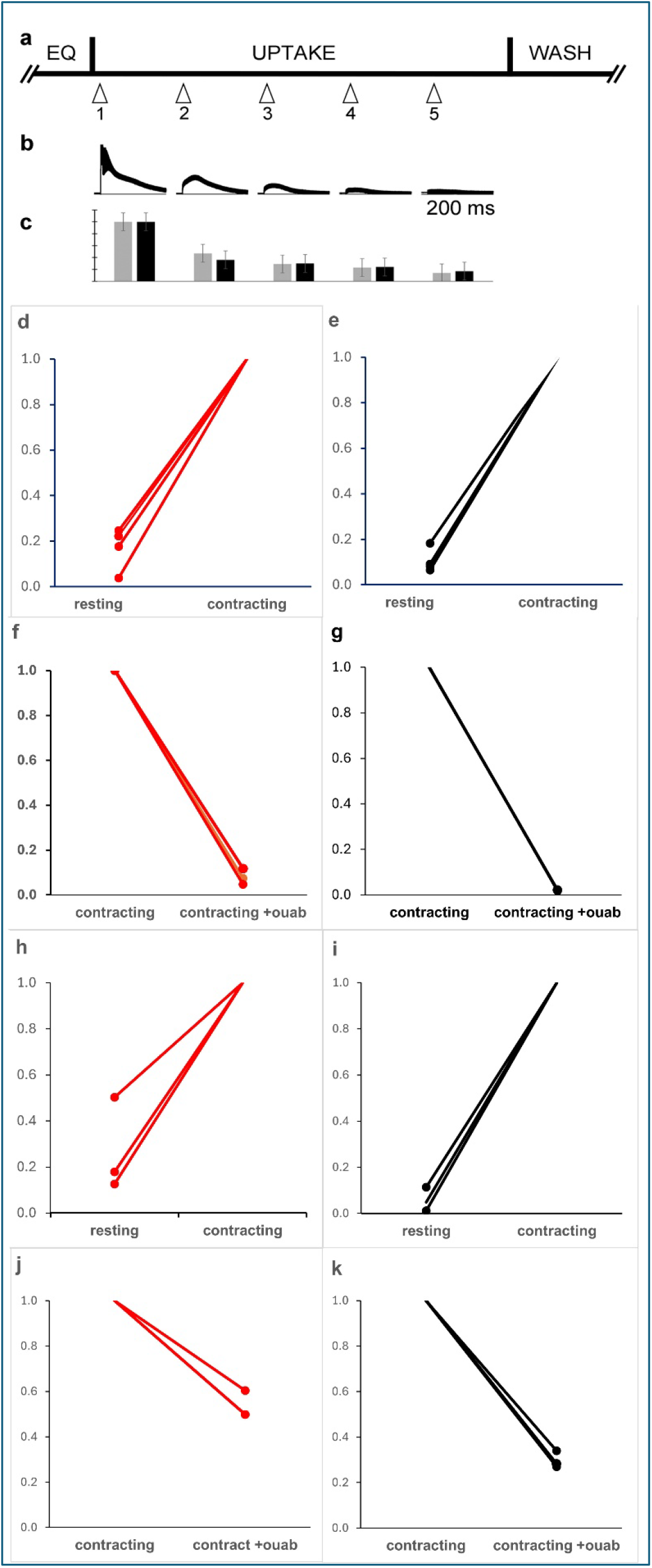
**a** Protocol to measure simultaneous uptake of Rb and glucose in isolated mouse EDL muscles. The muscle is perfused for 30 min at 32 °C in a standard Krebs solution (2 mL/min). Then, the solution is exchanged for Uptake Solution containing 200 μM tracer RbCl and 11 mM of a ^13^C-substituted glucose analog. Uptake is measured over a 5 min period, during which the muscle is maintained at rest or stimulated to produce 5 tetanic contractions (90 Hz/10 sec), repeated once per min. Following the uptake period, the muscle is processed for measurement of Rb and ^13^C content by ICP-MS. **b** Representative force recordings for tetani 1−5. Peak force 179.2 mN. **c** Normalized peak tetanic force in Uptake Solution without (grey) and with 0.7 μM ouabain (black). **d & e** Uptake of Rb (red) and 6-^13^C-glucose (black) in paired resting and contracting EDL muscles. αMDG uptake rates in resting and contracting muscles: 30.2 ± 21.9 and176.5 ± 38.0 μMol/min, respectively. p=0.002, t=15.2; n=3. **f & g** Uptake of Rb and glucose in muscles contracting in the absence and presence of 0.75 μM ouabain. **h & i** Uptake of Rb (red) and 6-^13^C-αMDG (αMDG, black) in paired resting and contracting EDL muscles. Rb, resting average: 42.8 ±1.6 nMol/min ; contracting average: 202.5 ± 33.1 nMol/min p=0.007, t=8.29, n=3; αMDG uptake rates in resting and contracting muscles: 30.2 ± 21.9 and176.5 ± 38.0 μMol/min, respectively. p=0.002, t=15.2; n=3. **j & k** Uptake of Rb and αMDG in muscles contracting in the absence and presence of 0.75 μM ouabain. Ouabain decreased Rb and αMDG uptake by 92 and 98 %, respectively; Rb, p=0.002, t=16.2; ^13^C, p=0.007, t=8.5; n=3. Ouabain decreased Rb and αMDG uptake by 92 and 98 %, respectively; Rb, p=0.002, t=16.2; ^13^C, p=0.007, t=8.5; n=3.

Resting muscles took up Rb and glucose at low rates (Fig. 1d,1e). These low basal uptake rates are expected because non-contracting muscles utilize only 3−5% of their total capacity for Na,K-ATPase transport ^18^, and basal glucose uptake is low in the absence of insulin. Muscle contraction dramatically stimulated uptake of both Rb and glucose (Fig. 1d, 1e) ^18,25^. Ouabain reduced Rb uptake by the Na,K-ATPase, as expected. Surprisingly, ouabain also reduced a fraction of glucose uptake (Fig. 1f, 1g).

Ouabain is not expected to inhibit glucose uptake by GLUT4, which uses a facilitated diffusion transport mechanism. This finding suggested that contraction-related glucose uptake may utilize, in part, an ATP-dependent glucose transporter.

To examine this possibility, we measured uptake of aMDG, a glucose derivative that is a substrate for SGLTs, but not GLUTs. Contracting muscles were able to take up αMDG (Fig. 1h,1i), which was completely inhibited by ouabain (Fig. 1j,1k), suggesting that the ouabain-sensitive component of glucose uptake may be mediated by an SGLT-type glucose transporter. The finding that inhibiting Na,K-ATPase-α2 transport induces a corresponding decrease in sugar uptake suggests that Na,K-ATPase-α2 transport is required for αMDG uptake.

To further examine the relationship between Rb and αMDG uptake, we stimulated Na,K-ATPase using the β_2_-adrenergic agonist, salbutamol (Fig. 2). Salbutamol stimulates Na,K-ATPase-α2 transport by approximately 2-fold in mouse EDL muscle ^24^ and lowers intracellular Na^+^. When NKA was stimulated by salbutamol, αMDG uptake increased in parallel (Rb 1.87-fold; αMDG 2.74-fold). Therefore, stimulation of Na,K-ATPase transport *per se*, independent of contraction, can induce αMDG uptake. Together, the ability to transport αMDG and the requirement for Na,K-ATPase active transport suggest that an SGLT contributes to insulin-independent glucose uptake in muscle.

**Fig. 2.**
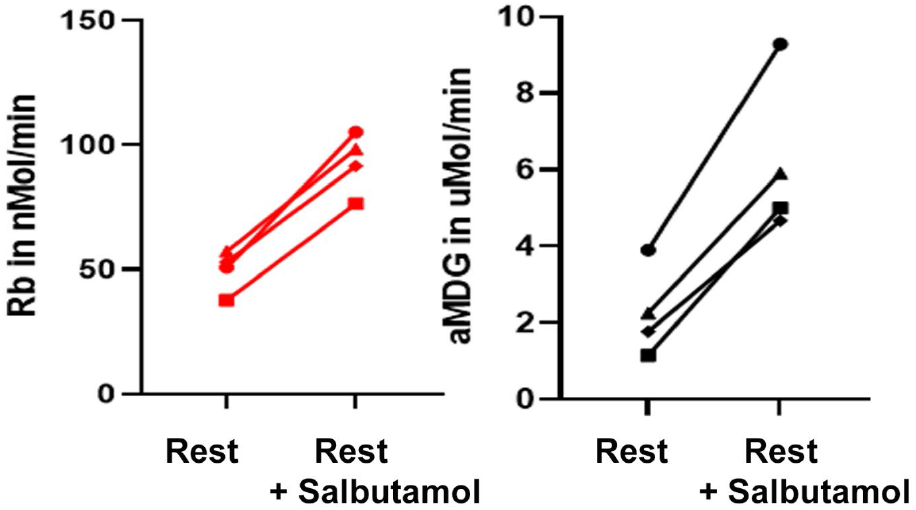
Rb and αMDG uptake in non-contracting EDL muscles in the absence and presence of 5 μM salbutamol. Rb: 49.8 ± 8.5; salbutamol: 92.9 ± 12.3, p=0.007 t=11.4; αMDG, rest: 2.3 ± 1.8; salbutamol: 6.2 ± 2.1 p=.002, t=7.6; n=4 EDL pairs..

### The αMDG transporter is distinct from GLUT4

To further distinguish mSGLT from GLUT4, we compared the effects of insulin and ouabain on uptake of 2DG and αMDG (Fig. 3) in non-contracting EDL muscles. Uptake of 2DG, stimulation by insulin, and lack of a requirement for Na,K-ATPase activity are hallmarks of GLUT4 transport ^29,30^. Other GLUT subtypes and SGLTs are not regulated by insulin, and most SGLTs have negligible affinity for 2DG as a substrate ^15^.

**Fig. 3.**
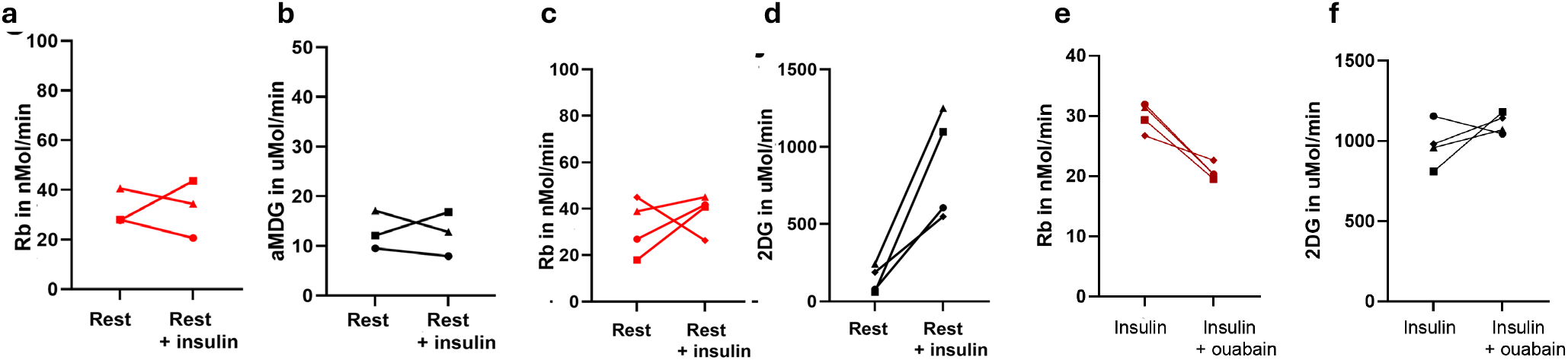
Uptake of αMDG and 2DG show different responses to insulin and ouabain. Rb (red), 2DG and αMDG (black). **a & b** Rb and αMDG uptake are insensitive to insulin (100 nM). Rb, rest: 32.1 ± 6.0; insulin: 32.8 ± 9.4 nMol/min, p=0.47, t=0.096; αMDG, rest: 12.9 ± 3.2; insulin: 2.5 ± 3.6 μMol/min, p=0.45, t=0.144; n = 3 EDL pairs. **c & d** uptake of 2DG, a hexose readily transported by GLUT4, is stimulated by insulin (positive control) without change in Rb uptake. Rb, rest: 32.1 ± 10.5; insulin: 40.9 ± 3.1 nMol/min, p=0.27, t=0.70; 2DG, rest: 141.2 ± 75.0; insulin: 874.5 ± 303.2 μMol/min, p=0.01, t=4.314 ; n=4 EDL pairs. **e & f** Rb and 2DG uptake in the presence of insulin ± 0.75 μM ouabain. Insulin-stimulated 2DG uptake is not inhibited by 0.75 μM ouabain (negative control), while ouabain decreased Rb uptake by 25%. Rb: 29.8 ± 2.4 and 20.7 ± 1.3 nMol/min, p= .013; 2DG: 975.6 ± 141.4 and 1109.4 ± 63.5 μMol/min, p = 0.27, n=4 EDL pairs.

Insulin did not alter Rb uptake (Fig. 3a, 3c), as expected. Insulin did not change αMDG uptake (Fig. 3b) but dramatically stimulated 2DG uptake (Fig. 3d). Insulin-stimulated 2DG uptake was not inhibited by ouabain (Fig. 3f). Ouabain at this concentration reduced Rb uptake by about 25% (Fig. 3e), indicating a modest effect on Na,K-ATPase-α2 activity. Taken together, the lack of response to insulin and ouabain exclude GLUT4 as the glucose transporter that mediates αMDG uptake.

### mSGLT uptake is additive with other glucose transport pathways

The data in Figs. 1−2 demonstrate that skeletal muscles can import glucose by a novel mechanism that transports αMDG, is not regulated by insulin, and can be stimulated by Na,K-ATPase-α2 activity. These properties are consistent with an SGLT-type glucose transporter and suggest that both mSGLT and GLUT4 are able to both import glucose during muscle contraction. To directly address this question, we measured uptake of 2DG and αMDG in paired EDL muscles during and post-contraction (Fig. 4). Muscle contraction stimulated Rb (Fig. 4a) and 2DG uptake (Fig. 4b), as expected. In this cohort, αMDG uptake was 16% of 2DG uptake during contraction (Fig. 4d) and 13% of 2DG uptake post-contraction (4f). Rb transport by Na,K-ATPase was independent of the hexose substrate (Fig. 4c, 4e), as expected since pump transport does not depend on the nature of the hexose. These data demonstrate that both GLUT4 and SGLT are able to take up glucose during and after contraction.

**Fig. 4.**
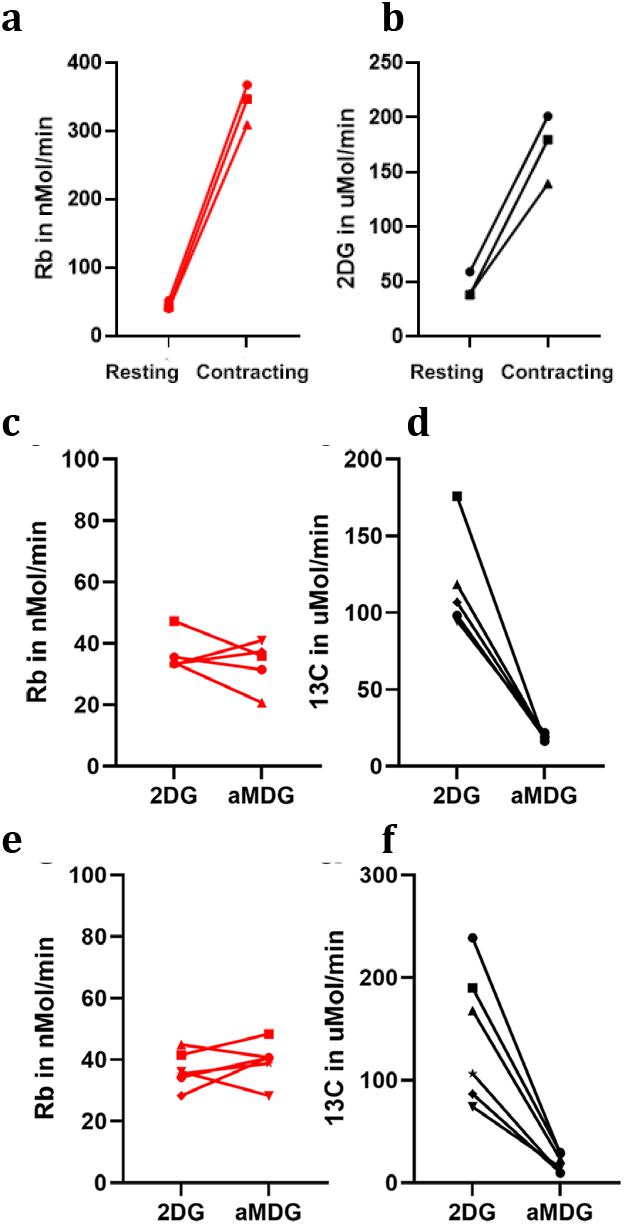
2DG and αMDG uptake both contribute during and following muscle contraction. **a & b** Muscle contraction stimulates Rb (red) and 2DG (black) uptake. Rb: p=0.001, t=21.1; 2DG: p=0.006, t=9.3; n=3 EDL pairs). **c & d** Rb and ^13^C uptake measured during contraction using either 11 mM 2DG or 11 mM αMDG as the hexose substrate. Rb uptake did not depend on the hexose substrate: p > 0.20, t=0.82, n=5 EDL pairs. **e & f** Rb and ^13^C uptake measured for 5 min post contraction. 2DG: 119.0 ± 33.1 μMol/min; αMDG: 19.0 ± 1.9 μMol/min, n=6 EDL pairs. Rb uptake did not depend on the hexose substrate: p

### Mouse skeletal muscle does not express a full-length SGLT2 transcript

Transcripts and short sequences of *Slc5a1* and *Slc5a2* genes, which contain 14 exons and encode SGLT1 and SGLT2, respectively, have been reported for human and mouse skeletal muscle using various expression methods [NCBI databases]. Here, we examined *Slc5a1* and *Slc5a2* expression using RT-qPCR. We detected *Slc5a2* mRNA sequences in mouse skeletal muscles but at low abundance. *Slc5a1* and *Slc5a2* sequences were detected in all replicates with 3 independent primer sets and 3 independent muscle samples (27/27 assays) with Ct values of 34-37. Given these high Ct values, we cannot exclude the possibility that the signal originates in non-muscle cell types within the whole muscle.

To further compare muscle with kidney transcripts, we amplified cDNA of *Slc5a2* from muscle and kidney using exon-scanning primer sets that target exons 2 – 12 (Table S3). We also amplified a near full-length *Slc5a2* coding transcript (Fig. 5). In kidney (Fig. 5a), amplicons of the expected size were detected for all region-specific primers (lanes 2−7) and a near full-length transcript (lane 8, 1896 bp of the 2070 bp coding sequence, consistent with reports that mouse kidney expresses a full-length transcript of *Slc5a2*. In muscle, none of the region-specific targets (lanes 10-15) nor a full-length transcript (lane 16) were detected on cDNA gel. The consistent low-level detection of short transcripts of *Slc5a2* in skeletal muscle (NCBI databases and this study) and absence of a full-length transcript of *Slc5a2*, suggest that mSGLT is distinct from the kidney SGLT2.

**Fig. 5.**
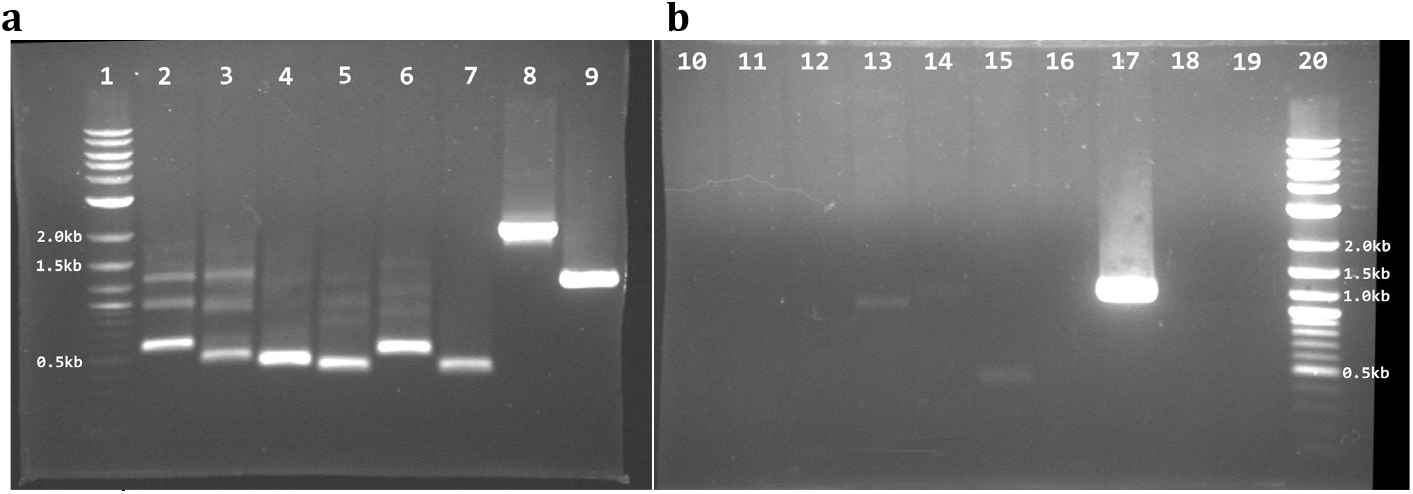
A near full-length transcript of *Slc5a2* is detected in mouse kidney (a, lane 8) but is absent in mouse skeletal muscle (b, lane 16). 889 ng of DNAse-treated kidney RNA (**a**) and 927 ng of DNAse-treated muscle RNA (**b**) was reverse transcribed, the cDNA amplified by PCR, and electrophoresed on a 1% agarose gel. Lanes 1 & 20, 1 kb+ ladder; Lanes 2−7, kidney RNA with primers that target regions of *Slc5a2* (primer sets 1−6, Table S3). Lane 8, kidney RNA amplified with a near full-length primer set (1896 bp of the 2070 bp coding sequence); lane 9, GAPDH (positive control); lanes 10−15, muscle RNA amplified with the same region-specific primers used for kidney; lane 16, muscle RNA amplified with the same full-length primer set used for kidney; lane 17, GAPDH ; lane 18, kidney no-RT control with GAPDH primers; lane 19, muscle no-RT control with GAPDH primers. cDNA from mouse kidney and hindlimb skeletal muscle was amplified using primer sets 1−6, 8, and GAPDH (Table S3). The same amount of cDNA was used as template for PCR of kidney and muscle. Gel is representative of 4 independent assays. The faint band in lane 15 (muscle, primer set 6 within exon 14), may be signal from the ubiquitous gene *Rusf1* which contains sequences shared with exon 14 of *Slc5a2*. For all primer sets, the melt curve peaks had identical melt curve/Tm values in kidney and muscle.

### Mouse skeletal muscle expresses a protein recognized by a SGLT2-specific antibody

In a further attempt to identify the muscle transporter, we performed immuno-histochemistry and Western blot assays using an SGLT2-specific antibody (ProteinTech 24654-1-AP, Fig. 6). The specificity of this antibody has been confirmed in kidney tissue from two independent *Slc5a2* knockout mouse models ^32,33^. This antibody targets a relatively less-conserved protein sequence among SGLT transporters (aa 180–311 of human SGLT2, corresponding to murine aa 178–309 (immunogen Ag19163, ProteinTech) and includes Ser287, an amino acid that is critical for αMDG binding ^34^. This SGLT2 antibody recognizes a protein in mouse muscle that co-localizes with Na,K-ATPase-α2 (Fig. 6a, 6b, 6c) in the muscle t-tubule membranes ^35^. In cross section, it labels fiber perimeters in a pattern typical of Na,K-ATPase-α2 in mouse fast muscle ^22^. The SGLT2 antibody colocalizes with MAP17, a subunit of SGLT2 that enhances activity ^34,36^. In longitudinal view (Fig. 6d, 6e, 6f), MAP17 and SGLT2 show a striation pattern typical of t-tubules, with two rows of t-tubules per sarcomere. In soleus muscle, a Myosin Type I antibody co-localizes with the SGLT2 label, confirming that the muscle αMDG transporter is expressed in both fast and slow fiber types (Fig. 56, 6h, 6i). Finally, the same MAP17 and SGLT2 antibodies label sections of human skeletal muscle (Fig. 6j, 6k, 6l), suggesting that a related protein is present in skeletal muscles of both species.

**Fig. 6.**
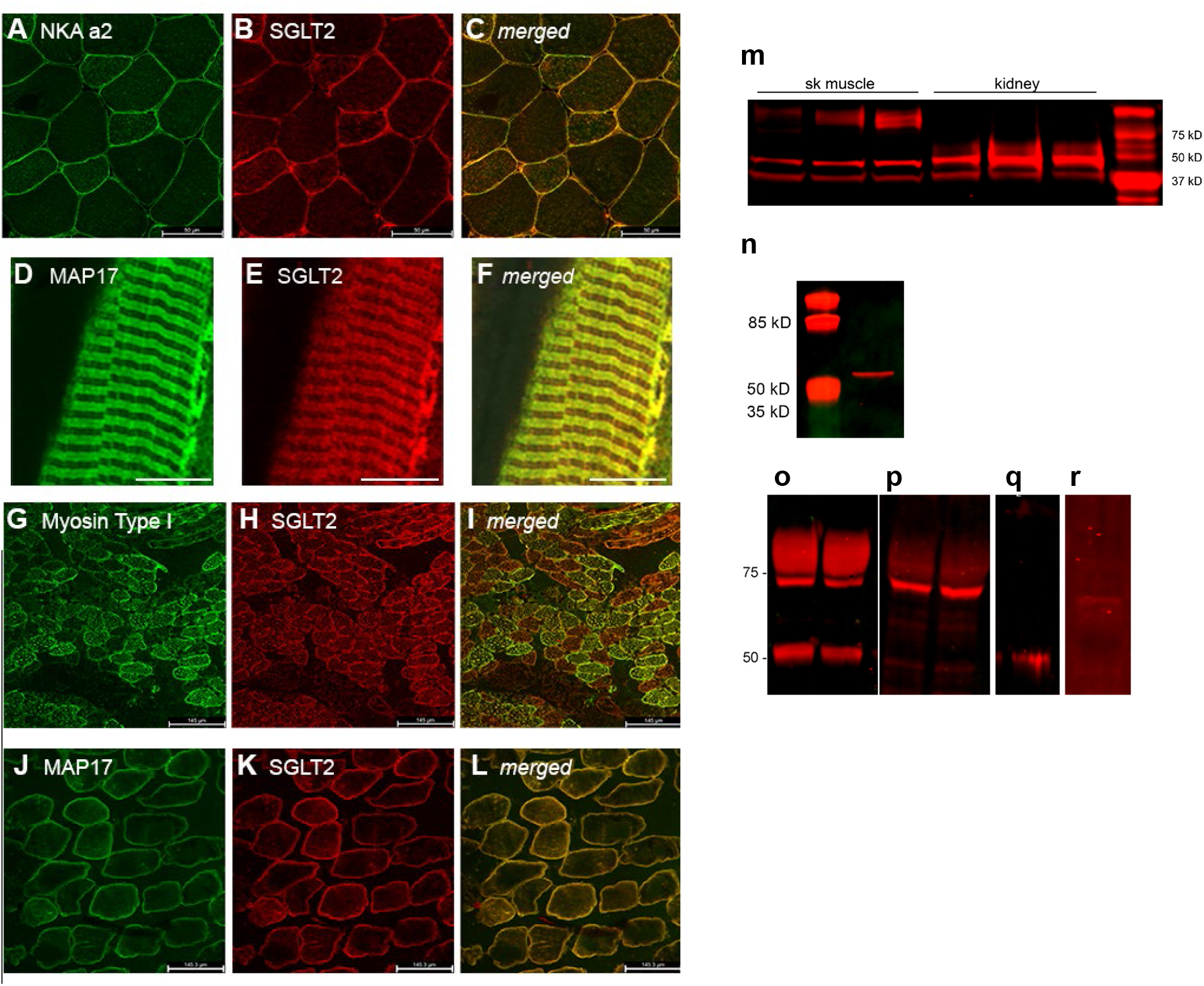
Immunodetection of SGLT2, MAP 17, and Na,K-ATPase-α2 in mouse and human skeletal muscle. **a – c** Confocal sections of fresh-frozen mouse tibialis anterior muscle dual-labeled with primary antibodies against Na,K-ATPase-α2 (green) and SGLT2 (red), and merged image. Scale bar 50 μm. **d – f** Longitudinal section of paraformaldehyde-fixed EDL fibers dual-labeled with MAP17 (green) and SGLT2 (red) antibodies, and merged image. Scale bar 13.5 μm. **g – i** cross-sections of mouse soleus muscle dual-labeled with myosin type I (green) and SGLT2 (red) antibodies, and merged image. Scale bar 145 μm. **j – l** Human vastus lateralis muscle dual-labeled with primary antibodies against MAP17 (green) and SGL2 (red), and merged image. Scale bar 145 μm. 10 μm sections. Images are representative of a minimum of three sections from each tissue sample. Negative controls included the secondary antibody and no primary antibody, and showed no signal for all sections (data not shown). Western Blot of mouse hindlimb skeletal muscle lysate (140 μg per lane) and mouse whole kidney lysate (20 μg per lane), labeled with the same SGLT2 antibody. **n** Western Blot of plasma membranes purified from mouse hindlimb skeletal muscle labeled with the same antibody. Muscle and kidney lysates were solubilized and denatured using a standard loading buffer with 2.5% SDS, 50 μM β-ME, and heated at 37 C for 30 min. **o, p** Western blots of purified plasma membranes from mouse hindlimb skeletal muscle (40 μg per lane) labeled with the same antibody used for IHC (PT) or an independent SGLT2 antibody (Abcam). The same muscle preparation was treated identically up to the antibody incubation step. This membrane preparation was solubilized and denatured more aggressively using 5% SDS, 100 mM DTT and heating at 65C for 15 min. **q, r** positive controls: human kidney cortical epithelial cells labeled with PT or Abcam antibodies. Western Blot images are representative of 3-4 blots obtained using 3 independent kidney and muscle samples. Control images without primary antibody and after Ponceau staining are provided in Extended Fig. S2. Western blot images are representative of 4 blots obtained using 3 independent kidney and muscle samples.

On Western blot, the same SGLT2 antibody recognizes a protein of ∼50 kD in whole lysates of mouse kidney and skeletal muscle (Fig. 6m). This is the size reported by this antibody in mouse kidney lysates and which was absent in kidneys of *Slc5a2* KO mice ^32,33^. It is found in purified plasma membranes (Fig. 6n), as expected for a membrane transporter and consistent with its membrane localization on immunohistochemistry. The size of the ∼ 50 kD band is less than the expected 72-75 kD for the full-length kidney SGLT2 (672 aa, 72 -75 kD depending on glycosylation); however, bands of either 50-55 kD or 70-75 kD have been reported for kidney SGLT2 using different antibodies and conditions. ^32,33,37^

We found that these differences are due, in part, to incomplete denaturation of the protein. When we denatured the samples using dithiothreitol instead of β-mercaptoethanol, we detected a protein of ∼ 72 kD in both muscle and kidney. This band is also detected by an independent SGLT2 antibody (Abcam ab37296, Fig. 6o, 6p). DTT is a 7-fold more potent denaturant than β-mercaptoethanol and may be needed for a protein as hydrophobic as SGLT2, which has 14 transmembrane helices and multiple disulfide bonds.

In summary, two independent SGLT2-specific antibodies detect a protein in muscle that has the same size as the kidney SGLT2, and its migration pattern on gels tracks the apparent size of kidney SGLT2 under different denaturing and solubilization conditions. The muscle protein shows a plasma membrane localization in immunohistochemistry and Western blot. These concordances demonstrate that skeletal muscles express a protein that shares antigenic regions with the kidney SGLT2.

### Phlorizin and empagliflozin show weak inhibition of mSGLT uptake

We also attempted to identify the muscle transporter using pharmacological inhibitors of SGLT transport (Fig. 7). The affinities of SGLTs for hexoses and competitive inhibitors is a distinguishing feature of SGLT subtypes ^15^. In contracting muscles, 10 μM phlorizin did not produce a consistent action on either Rb or αMDG uptake (Fig. 7a, 7b). The failure of phlorizin to inhibit uptake at concentrations much greater than its IC^50^ for SGLT1 (21 nM) or SGLT2 (290 nM) ^38^ excludes these subtypes as the transporter that mediates αMDG transport in muscle. However, interpretation of the action of phlorizin is not conclusive due to its known off-target actions on the Na,K-ATPase. Phlorizin can stimulate or inhibit enzyme activity ^39,40^ depending on K, Na and glucose concentrations. In our conditions, greater concentrations of phlorizin consistently inhibited the Na,K-ATPase and were not investigated further. Empagliflozin, 500 nM inhibited salbutamol-stimulated NKA activity but, paradoxically, increased αMDG uptake (Fig. 7c, 7d). In contracting muscles (Fig. 7e, 7f), empagliflozin did not alter Na,K-ATPase activity and decreased αMDG uptake by 25%. The failure of empagliflozin to inhibit αMDG uptake at a concentration far greater than its IC^50^ for SGLT2 (3 nM), excludes SGLT2 as the tranporter that mediates αMDG transport in muscle.

**Fig. 7.**
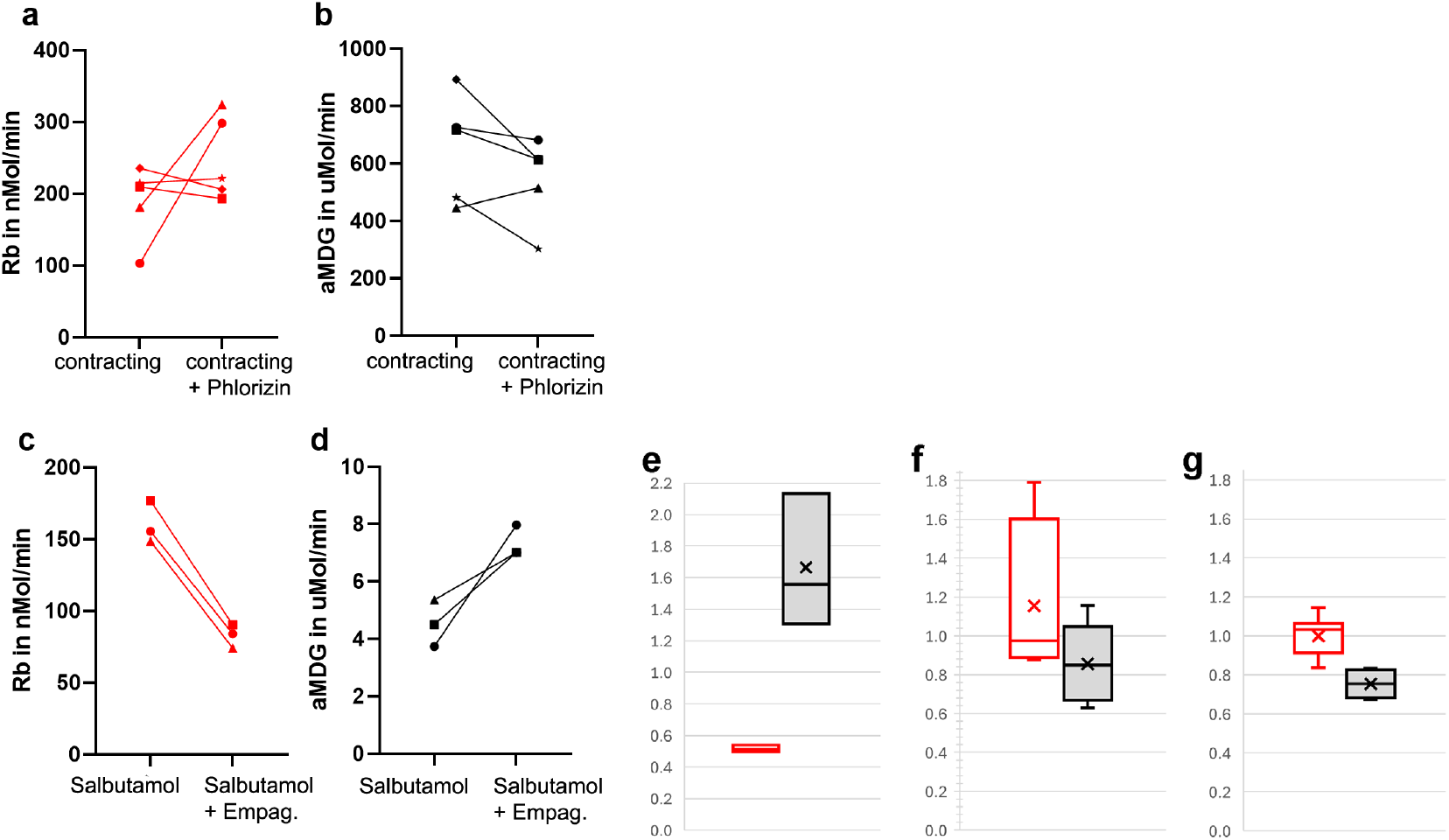
Inhibitors of SGLT1 and SGLT2 show weak affinity for mSGLT. **a & b** Uptake of Rb (red) and αMDG (black) in the absence or presence of 10 μM phlorizin. Rb: 244.0 ± 27.1 and 249.0 ± 58.0 nMol/min in control & phlorizin, respectively, p=0.82; αMDG: 5.9 ± 1.4 and 6.8 ± 0.8 μMol/min, p = 0.27. n= 5 EDL pairs. **c, d** EDL muscles incubated in 5 μM salbutamol to stimulate the Na,K-ATPase, in the absence or presence of 500 nM empagliflozin. Rb: 177.8 ± 32.0 and 88.7 ± 10.8 nMol/min in control and empagliflozin, respectively; p = .006; αMDG: 5.4 ± 1.6 and 6.9 ± 0.9 μMol/min in control and empagliflozin; p = 0.19, n=3 pairs. **e** EDL muscles contracting without and with 10 μM phlorizin. Ratio uptake in drug/uptake in vehicle. Rb (red) 1.15 ± 0.43 (5). αMDG (black) 0.86±0.21 (5); **f** EDL muscles stimulated with salbutamol without or with 500 nM empagliflozin. Rb (red) 0.52 ± 0.02 (3). αMDG (black) 1.67±0.42 (3). **g** EDL muscles contracting without and with 500 nM empagliflozin. Rb (red) 1.00 ± 0.10 (9). αMDG (black) 0.75±0.07 (4).

## DISCUSSION

### Summary

This study demonstrates that a previously unrecognized glucose transport pathway contributes to non-insulin mediated glucose uptake in skeletal muscle. This uptake pathway is not regulated by insulin, requires Na,K-ATPase-α2 activity, and transports the hexose αMDG. Because these are defining properties of SGLT type glucose transporters, we refer to it as “mSGLT”.

Our results suggest a functional coupling between Na,K-ATPase-α2 and mSGLT transport in the muscle t-tubules, the major site of glucose disposal into muscle. Stimulation of Na,K-ATPase-α2 activity by contraction or the catecholamine, salbutamol, induces mSGLT transport, while inhibition of Na,K-ATPase-α2 activity with sub-micromolar ouabain inhibits it. We propose a model in which an SGLT-type glucose transporter in the muscle t-tubules imports glucose together with Na^+^ using the inward Na gradient generated by Na,K-ATPase-α2 (Fig. 8). This pathway is independent of and additive with the non-insulin triggered GLUT4 pathway. (insulin-triggered GLUT4 uptake is absent in our insulin-free conditions).

**Fig. 8.**
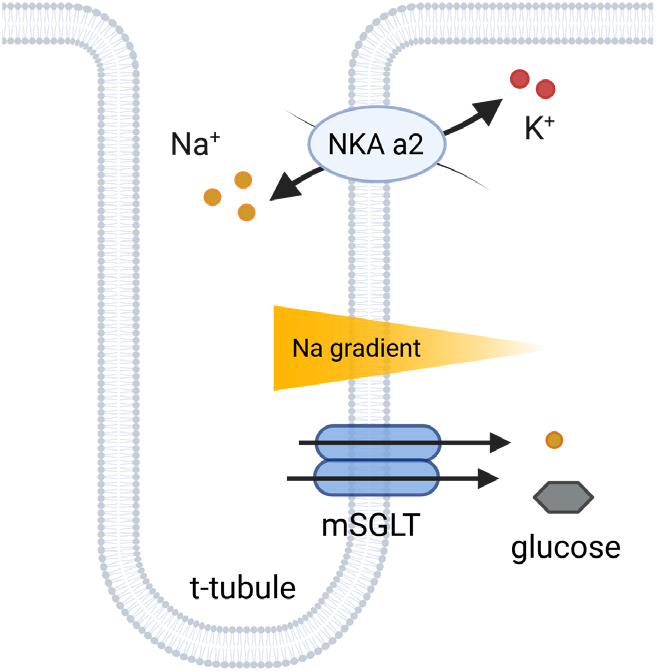
Proposed model of mSGLT transport. A muscle sodium-glucose linked transporter, mSGLT, imports glucose together with Na^+^. mSGLT uses the inward Na gradient generated by Na,K-ATPase-α2 transport to drive Na^+^ into the muscle, and glucose follows along with Na^+^. Both transporters localize to the t-tubule membranes. By providing the inward Na gradient, the Na,K-ATPase-α2 makes an indirect contribution to sodium-glucose linked transport by mSGLT.

The ATP that fuels Na,K-ATPase-α2 transport comes largely from glycolysis and is generated locally by glycolytic enzymes in the t-tubule membranes ^41^. The physical positioning in the same membrane of gl^42^ucose transporters to provide energy substrate, glycolytic enzymes, and a major ATP consumer (Na,K-ATPase-α2) provides an efficient mechanism to support muscle contraction.

Our discovery of this pathway was enabled by the technology we developed to measure simultaneous uptake of co-transported species, including both inorganic ions and carbon-based organic substrates. ^25^. Most previous investigations of glucose uptake in muscle have measured uptake of a single, radiolabeled glucose or glucose derivative that is a substrate for GLUTs. To our knowledge, neither Na,K-ATPase substrates nor αMDG, a substrate not handled by GLUT4, have been examined previously in mouse muscle, so this pathway would not have been seen. Prior studies of glucose uptake into muscle largely measured glucose uptake following contraction, due to technical limitations of radioisotope use.

With our method, we were able to measure glucose uptake in intact muscles under basal, contracting, and post-contraction conditions. Muscle metabolism differs greatly in these states, as muscles shift from glucose utilization to glucose replenishment and storage. The presence of glucose transporters having different transport mechanisms and regulation may allow muscles to match glucose uptake to a wider range of physiological conditions than possible with GLUT4 alone. GLUT4 uses a translocation mechanism and the inward glucose gradient to import glucose by facilitated diffusion; while, SGLTs use the inward Na gradient generated by Na,K-ATPase activity to import glucose together with Na^+^ ^43^. Both gradients vary dynamically during and following muscle contraction and in different metabolic conditions. Interestingly, maximum insulin stimulation (100 nM) or electrical stimulation of isolated muscles each induces a similar, 2-3-fold increase in the surface density of GLUT4, suggesting that GLUT4 translocation is saturable ^42,44 45^. The presence of a glucose transporter driven by the Na gradient may allow muscles to continue to function under conditions when GLUT4 translocation is saturated or impaired.

#### Contribution of mSGLT to non-insulin mediated glucose uptake into muscle

In this study, the magnitude of contraction-stimulated αMDG uptake by mSGLT varied in different experimental cohorts. The variability tended to correlate with tetanic force and degree of fatigue, suggesting that initial energy reserves or glycogen stores may influence its contribution to non-insulin mediated glucose uptake. In addition, the transmembrane gradients for Na^+^ and glucose, which drive SGLT and GLUT4 transport, respectively, vary dynamically during contraction. The relative amounts of 2DG and aMDG uptake measured in the conditions of Fig. 3 cannot be taken to indicate the contributions of GLUT4 and mSGLT to total glucose uptake under physiological conditions. This determination requires knowledge of the substrate profile of mSGLT and its substrate affinities for αMDG relative to glucose, the natural substrate in vivo. This characterization is best done using an expression system or other well-controlled model after mSGLT has been identified and sequenced. Although the best-characterized SGLTs have only a low affinity for 2DG, the affinity of mSGLT for 2DG is not known. If mSGLT has a broad substrate affinity, the measured 2DG uptake may report contributions from both GLUT4 and mSGLT. It will be important in future studies to define the relevant parameters that determine their separate contributions.

### Physiological significance

The skeletal muscles play a major role in glucose homeostasis due to their large capacity to take up, store, and utilize glucose. Muscles need both insulin-mediated and non-insulin-mediated pathways to achieve glucose homeostasis. Non-insulin mediated glucose uptake operates during periods of declining insulin and elevated catecholamines, conditions that occur acutely during exercise and chronically during periods of stress, after injury, and in metabolic disorders^46^. Epinephrine and norepinephrine released by contracting muscles or during stress inhibit pancreatic insulin release and reduce the insulin sensitivity of muscle. Importantly, non-insulin mediated glucose uptake is preserved in obesity and diabetes, when it contributes 70% of total glucose disposal ^47^. In these conditions, the presence of a non-insulin-dependent glucose transporter with muscle-specific expression may be essential for normalizing blood sugar.

### Molecular identity of mSGLT

SGLT subtype mRNA transcripts have been reported previously in skeletal muscles of human, rat, and mouse and attributed to SGLT2 expression. ^48-50^ However, detection of short transcripts in a gene family with high homology does not indicate that the protein is expressed and functional. We also consistently detected short RNA sequences spanning exons 4-11 of *Slc5a2* and exons 3-12 of *Slc5a1* by RT-qPCR. However, their copy numbers were very low (Ct > 30) compared to kidney (Ct= 24), and a full-length transcript of *Slc5a2* was not present. Low-level detection may suggest a low abundance transcript, weak hybridization with SGLT2 primers, the presence of a transcript variant of SGLT2 or related gene, or a contribution from non-muscle cells within the tissue. Our finding is consistent with a previous study in which SGLT2 mRNA was detected in kidney (Ct = 24) but below the level of detection (Ct >30) in mouse skeletal muscle ^37^. Taken together, these findings argue that *Slc5a2*/SGLT2 is not expressed in mouse skeletal muscle.

This conclusion is further supported at the protein level. A SGLT2-specific antibody validated in independent knockout models ^32,33^ labeled a protein in the plasma membranes with both immunohistochemistry and Western blot assays, as expected for a membrane transporter. On Western blot, two independent SGLT2 antibodies label the same two proteins reported in kidney -a 72 kD protein near the expected size of the full-length SGLT2 and a protein of ∼ 55 kD. Both proteins are products of the *Slc5a2* gene, as validated in three independent knockout models ^32,33,37^. The 50 kD protein may result from post-transcriptional or post-translational processing of *Slc5a2*, or may be cleavage in the assay conditions. These results indicate that the muscle transporter, mSGLT, shares antigenicity with the kidney SGLT2. These concordances demonstrate that skeletal muscles express a protein that shares antigenic regions with the kidney SGLT2. However, antibody recognition alone cannot be used to conclude that the muscle protein is identical with the kidney SGLT2 because skeletal muscle does not express a full-length SGLT2 mRNA transcript. In addition, these antibodies were generated using immungens that target short sequences of the kidney SGLT2. They were optimized to distinguish SGLT2 from SGLT1, but not other possible SGLT isoforms.

Pharmacological tests with the competitive inhibitors phlorizin and empagliflozin further exclude SGLT1 and SGLT2 as candidate transporters for mSGLT. A concentration of empagliflozin more than 160-times its IC_50_ for SGLT2 inhibited αMDG uptake by only 25%, indicating that mSGLT is a low affinity substrate for empagliflozin.

Taken together, our gene, protein, and pharmacological evidence suggest that mouse skeletal muscles express a protein that shares antigenic regions and some transcript homology with SGLT2/Slc5a2 but is not identical with kidney SGLT2. mSGLT may be produced by a transcript variant of *Slc5a2* in muscle or may be the product of a different gene. Na-linked transporters comprise a large gene family with a high degree of homology. A complete molecular identification of mSGLT will require additional approaches, including sequencing and proteomics.

## METHODS

### Tissue sources

Adult male and female C57BL/6 mice (The Jackson Laboratory or Charles River Laboratories) at 2−3 months of age were used as a source of tissue. All procedures involving animals accorded with the *Guide for the Care and Use of Laboratory Animals* (National Research Council of the National Academies, USA) and were approved by the University of Cincinnati Institutional Animal Care and Use Committee. Mice were housed in a 12:12 hour light-dark cycle in standard rodent cages with free access to standard rodent chow. For tissue extraction, mice were fasted 3 h prior, anesthetized (2.5% Avertin, 17 ml/kg), and euthanized after tissue removal. Frozen human vastus lateralis muscle samples were obtained from the University of Kentucky Center for Muscle Biology. The human kidney cell line, TERT1-HK2, was kindly provided by Dr. Czyzyk-Krzeska, University of Cincinnati.

### Chemicals

Chemicals were sourced as follows: ouabain (Sigma-Aldrich); 6-^13^C-2-deoxy-D-glucose, (CAS 201740-77-4, Cambridge Isotope Laboratories, Inc.); human insulin (CAS 11061-68-0, Sigma-Aldrich); phlorizin (CAS 60-81-1, Tocris Bioscience; and CAS 7061-54-3, Sigma-Aldrich); empagliflozin (S8022, Selleck Chemicals) 6-^13^C-alpha-methyl-D-glucoside (CAS 97-30-3) was custom synthesized (laboratory of W. Priebe, MD Anderson, TX); Certified Reference Material (CRM) trace metal drinking water (CRM-TMDW, High-Purity Standards, USA; CRM milk powder (CRM-MP, High-Purity Standards, USA); inorganic carbon standard (Inorganic Ventures, USA); bovine liver CRM (NIST #1577b); phosphorous CRM (NH_4_H_2_PO_4_ in H_2_O, High Purity Standards #IC-P-M); multi-elemental solution 2 (SPEX CertiPrep #CLMS-2N); river sediment CRM (High purity Standards #CRM-MS-S). All other chemicals were trace metal grade (Sigma Aldrich or Thermo Fisher Scientific).

### Experimental solutions

For isolated muscle experiments, the Equilibration Buffer contained (mM): 118 NaCl (Sigma), 4.7 KCl, 2.5 CaCl_2_, 1.2 MgCl_2_, 1.2 NaH_2_PO_4_, 11 D-glucose, 25 NaHCO_3_; gassed with 95% O_2_, 5% CO_2_; pH 7.4, 32 °C. The Uptake Buffer contained (mM): 118 NaCl, 4.7 KCl, 2.5 CaCl_2_, 1.2 MgCl_2_, 1.2 NaH_2_PO_4_,11 6-^13^C-2DG or 11 6-^13^C-αMDG [Extended Fig. S1], 25 NaHCO_3_; and 200 μM RbCl pH (7.4, 32 °C). The Na-free Wash Buffer contained (mM): 15 Tris-Cl, 2.5 CaCl_2_, 1.2 MgCl_2_, 263 sucrose; pH 7.4, 0–2 °C; or: 10 Tris-Cl, 2.8 KCl, 1.2 KH_2_PO_4_, 1.3 CaCl_2_, 1.2 MgCl_2_; pH 7.4, 32 °C. Solutions were prepared using 18 MΩ-cm water, metal-free containers or acid-washed (2% HNO_3_) glassware. Solutions were stored at 4 °C and used within one week of preparation.

### Measurement of glucose and Rb uptake

Uptake of Rb, ^13^C, and other ions was measured in isolated EDL muscles obtained from anesthetized female mice. Measurements were made under nominally insulin-free conditions to exclude glucose uptake by known insulin-dependent mechanisms. This was achieved by fasting the mice for 3 h prior to muscle extraction to reduce circulating insulin ^51^, using isolated muscles removed from the circulation, perfusing the muscle for 30 min prior to uptake measurement, and omitting insulin from all solutions. Two EDL muscles (test and control) and two tibialis anterior (TA) muscles were taken from each mouse. The TA muscle was untreated and assayed in parallel to obtain the endogenous Rb content of untreated muscles and the experimental ^13^C/^12^C ratio under no-^13^C exposure conditions. The endogenous Rb concentration varies 5−10% in different animals but is highly consistent (<2%) in different muscles from the same animals ^52^. The protocol for measuring sugar and Rb uptake was essentially as described ^25,52^.

The test EDL muscle was positioned between parallel plate electrodes in an experimental chamber and perfused with Equilibration Buffer at a flow rate of 2 ml/min. The temperature of the perfusate was maintained at 32 °C using an in-line heater and monitored by a bath thermistor positioned near the muscle. One tendon was fixed and the other tendon was attached to a force transducer. The muscle length was set to the optimal length, Lo, at which the muscle produces peak twitch force. After a 30 min perfusion with Equilibration Buffer, the solution was rapidly exchanged for Uptake Buffer containing 11 mM of a ^13^C-substituted glucose substrate and 200 μM RbCl, which served as a tracer for K transport by the Na,K-ATPase. The muscle was maintained in Uptake Buffer for 5 min, during which the muscle was either left at rest or stimulated electrically to produce repetitive tetanic contractions. At the end of the uptake period, the muscle was perfused with Wash Buffer at 0 °C to inhibit membrane transport, then removed from the chamber and washed extensively to remove excess cations from the extracellular space. The wash protocol consisted of four consecutive 5 min incubations in 10 mL Wash Buffer at 0 C with shaking. After washing, the muscle was gently blotted, weighed on an analytical balance, and stored at 4 °C until processed for measurement of the ^13^C, Rb, and P content by ICP-MS. Rb uptake was computed as the amount of Rb taken up by the muscle during the 5 min uptake period after subtracting the naturally abundant, endogenous [Rb] in the muscle (obtained from untreated TA muscles). Rb uptake was computed as the ouabain-sensitive component of Rb uptake, obtained using 0.75 μM ouabain in Equilibrium and Uptake solutions. In measurements using ouabain, phlorizin, or empagliflozin, the drug at the indicated concentration was included in both Equilibration and Uptake Buffers, to allow binding to reach steady state. The contralateral muscle of each mouse was subjected to the identical protocol but without the test manipulation and served as control. The order of test and contralateral measurements was randomized. Test and control EDL muscles together with a TA muscle from the same animal were processed identically in parallel for ICP-MS, as described below.

### Measurement of ^13^C-substituted hexoses, Rb, and P by ICP-MS

All steps were performed under metal-free conditions. Muscle samples were prepared for ICP-MS using a sequence of steps designed to remove background C from insoluble cellular components, largely membranes and proteins, and obtain a soluble cell fraction containing the target elements (^13^C, ^12^C, P, Rb). After the final wash, samples were stored at -20 °C for up to 5 h, then processed as follows: freeze in liquid N_2_ and lyophilize overnight; solubilize in LCMS Optima grade water (1.5 mL for EDL, 3 mL for TA) and homogenize using a motorized Potter-Elvehjem homogenizer, 20 rpm for 5 min; sonicate using 3 × 100 mm ultra-sonication probe at 37.5 watts, 2 sec pulses delivered every 5 sec for 1 min; centrifuge to pellet insoluble components (450 x g for 5 min, followed by 13,000 x g for 10 min; acidify with 5% v/v HCl to precipitate cytosolic proteins; freeze overnight at -20 °C; thaw and centrifuge 13,000 x g for 10 min. The final supernatant contained soluble cytosolic components including the target elements. This supernatant was used directly for C analysis by ICP-MS. For Rb analysis, an additional dilution (5x for EDL, 10x for TA) in 2% HNO_3_ together with internal standards (Sc and Y) was made. ICP-MS analysis was performed using tuning parameters (Extended Data Table S1). The instrument was tuned daily and carbon calibration curves were obtained as described ^25^. For P calibration, a 10 ppm working standard was made in HPLC Optima grade water and diluted serially to have 0, 0.05, 0.1, 0.5, 1, 2.5, and 3 ppm standards. For Rb calibration, a Rb working standard of 100 ppb in 2% HNO_3_ was used to make 0, 0.5, 1, 5, 10, 20, and 30 ppb samples. A multi-element working standard was used to make 0, 50, 100, 500, 1000, 2000, and 3000 ppb of Na, Ca, K, and Mg. In addition to the target elements, the P content of each muscle sample is measured and used as an accurate mass index for normalization ^24^. Uptake rates for hexose and Rb were computed from ICP-MS measurements as described ^25^, except using the P content of each muscle as a mass index for normalization rather than wet weight. Uptake rates were measured in ng/g, normalized to the P content of each muscle as a mass index, and converted to nMol/min as follows:

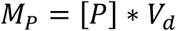

Where M_p_ is the total mass of P in nanograms, V_d_ is the final dilution volume of the samples after processing (31.5 mL for TA and 7.88 mL for EDL), and [P] is the concentration of P measured by ICP-MS.

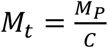

Where M_t_ is the mass of tissue in grams and C is a constant value equal to 2.5x10^6^ ng of P per gram of mouse tissue [2500000 ng/g dry tissue mass]

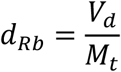

Where d_Rb_ is the dilution factor of Rb.

To compute the total concentration of Rb in the sample in ppb, multiply d_Rb_ with the concentration of Rb as measured by the ICP-MS. The amount of Rb taken up by the muscle during the uptake period is computed as the total [Rb] taken up by the EDL during the uptake period minus the endogenous concentration of Rb obtained from the tibialis anterior muscle of the same mouse. To get the concentration of Rb from ppb to the rate of Rb uptake in nMol/min, simply divide the concentration of Rb in EDL by the mass of Rb (85) to get nanomoles then divide by the total uptake time.

### Gene expression

Total RNA was isolated from mouse tissues using RNAzol-RT (Molecular Research Center, Inc., Cincinnati, OH). RNA pellets were dissolved and stored in FORMAzol (Molecular Research Center, Inc.) at -80 C. For subsequent reactions, RNA was precipitated from FORMAzol, resuspended in nuclease-free water, treated with DNase I (Thermo Fisher Scientific), and quantified (NanoDrop ND-2000C Spectrophotometer, Thermo Fisher Scientific). *Slc5a* gene family transcripts were measured by RT-qPCR (StepOnePlus Real-Time PCR System (Applied Biosystems); iTaq Universal Probes One-Step Kit (Bio-Rad)). Each sample was assayed in triplicate using 100 ng total RNA per reaction. Primers and probes used for Table 1 were selected from Integrated DNA Technologies pre-designed PrimeTime qPCR Probe Assays and screened using NCBI BLAST to confirm the lack of off-target hybridization. A standard curve using reference samples of pooled kidney RNA (20 ng, 4 ng, 0.8 ng, and 0.16 ng RNA) was run with each plate to ensure consistency and validate comparisons among different plates. Identification of specific sequences within *Slc5a2* was performed using PCR and gel electrophoresis. cDNA was generated with SuperScript IV First-Strand Synthesis System (Thermo Fisher Scientific) and Oligo d(T)20 primer. PCR was performed using an Eppendorf Mastercycler Pro with PrimeSTAR-HS (Premix) (Takara Bio) and primers at a final concentration of 0.25 μM each. All primers were from Integrated DNA Technologies. 10% of RT and no-RT reactions were used as template in 20 μl PCR reactions. PCR control reactions (water, no template) were included in all runs. PCR conditions were: denature at 98 °C for 10 seconds, anneal at 60 °C for 15 seconds, extend at 72 °C for 2 minutes, for a total of 30 cycles. The entire 20 μl of each DNA product was electrophoresed together with a 1 kb Plus DNA ladder (New England BioLabs) on a 1% TopVision Agarose gel (Thermo Fisher Scientific).

**Table 1.**
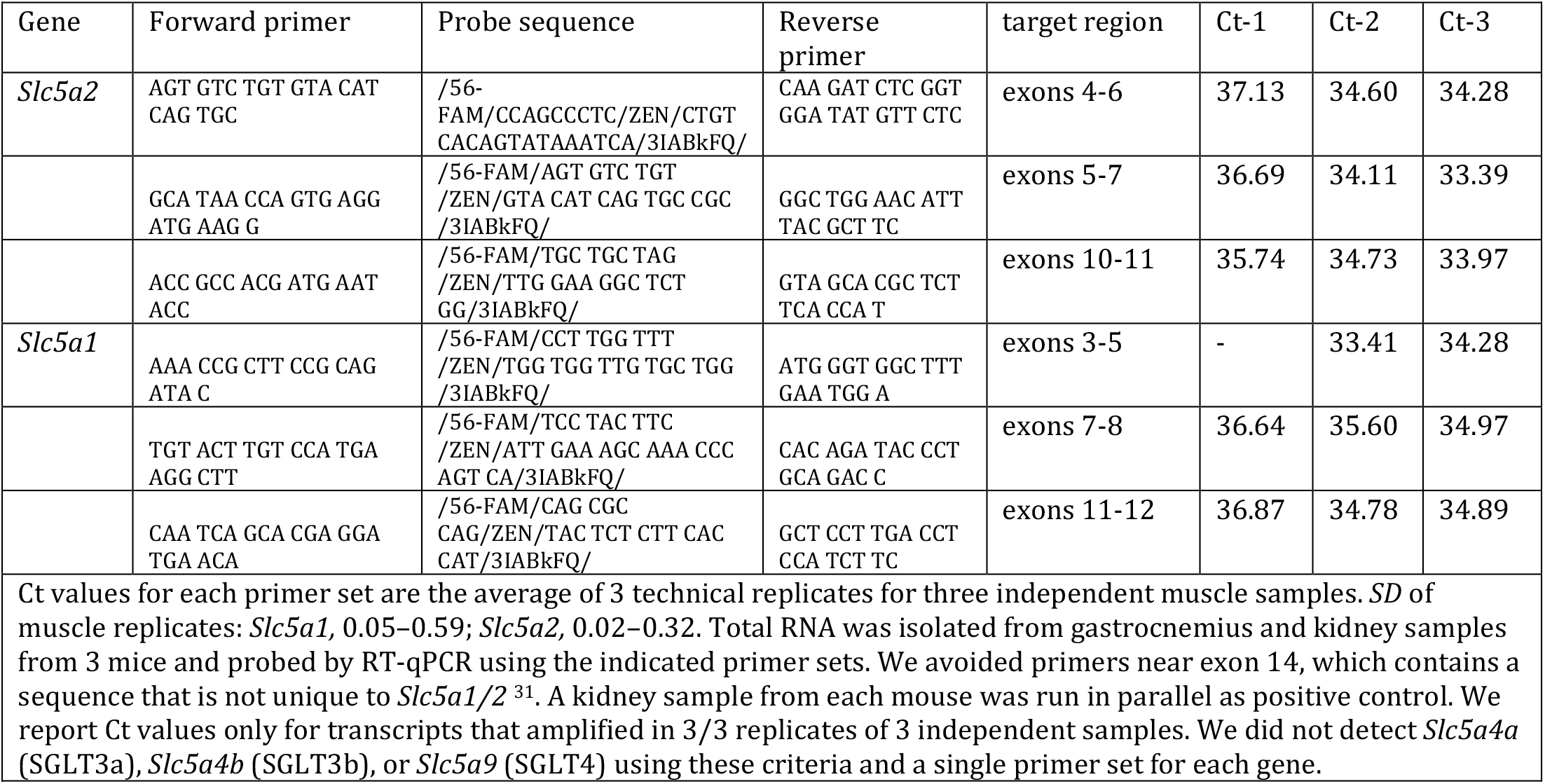
Transcripts of *Slc5a2* and *Slc5a1* are detected in mouse skeletal muscle.

### Immunohistochemistry

Tibialis anterior muscles were removed from anesthetized male and female mice at 8–16 weeks age. The muscles were coated with Tissue Plus O.C.T. compound (Fisher Health Care), snap frozen in liquid nitrogen, and stored at -80 °C until processed for image analysis. Transverse 10 μm sections were cut at -22 °C (Micron HM 525 cryostat), dried in air at room temperature for 2 min on microscope slides (Fisher Superfrost Plus Gold precleaned), fixed with 100% acetone at -20 °C, and transferred to Phosphate Buffered Saline (PBS). Permeabilization was performed at 65 °C in 2 mM sodium citrate with 0.5% Tween-20. Sections were incubated overnight at 4 °C in the primary antibody (1:50 in PBS), washed 3×7 min in room temperature PBS, incubated in the secondary antibody (1:200 in PBS) for 1.5 h at room temperature, washed 3×7 min in room temperature PBS, mounted with mounting media (Prolong Gold Antifade Mountant, refractive index 1.46; Thermo Fisher Scientific) and cover glass (0.13-0.17 mm thickness, FisherBrand 22×50). Sections processed identically but without primary antibody served as negative controls. Primary antibodies were rabbit anti-SGLT2 (24654-1-AP, ProteinTech ^53^), rabbit anti-Na,K-ATPase-α2 (16836-1-AP, ProteinTech), rabbit polyclonal anti MAP17 (PA5-53252, Invitrogen), mouse monoclonal anti-Myosin heavy chain (slow α and β, BA-F8-c, DSHB, University of Iowa). Secondary antibodies were goat anti-rabbit Alexa Fluor Plus 488 conjugate of F(ab’)_2_ fragment (Life Technologies), donkey anti-goat IRDye 680RD (Li-Cor Biosciences), and goat anti-rabbit IgG IRDye 800CW (Li-Cor Biosciences). Imaging of double-labeled sections was performed at room temperature using a Leica Stellaris 8 confocal microscope equipped with 63× oil-immersion objective and LAS-X acquisition software. Human muscle sections were processed as described for mouse muscle. For images of dissociated EDL fibers, paraformaldehyde fixed fibers were moderately stretched, pinned to cork, incubated in freshly made 1% paraformaldehyde in PBS pH 7.4 for 30 min at room temperature. Processing was the same as for sections up to the permeabilization step when bundles of 2−3 fibers were manually separated and mounted on glass slides.

### Western Blot

Hindlimb muscles were dissected from anaesthetized mice, frozen in liquid nitrogen, and homogenized in cold Lysis buffer (Pierce Lysis Buffer, Thermo Fisher) using a Tissue-Tearor (Bio-Spec Products, 3× 30 sec at max speed). To obtain total cell lysate (Fig. 6m), the homogenate was centrifuged at 4 °C for 10 min, at 3,300x g. To obtain plasma membrane proteins (Fig. 6n), the total cell lysate was purified using the Minute™ Plasma Membrane/Protein Isolation and Cell Fractionation Kit (Invent Biotechnologies) following the manufacturer’s protocol. To obtain a total membrane fraction from muscle (Fig. 6o), the cell lysate was centrifuged at 180,000xg for 60 min. For antibody positive controls, a cell lysate was obtained from whole mouse kidney (Fig. 6m) or from a human kidney cell line, TERT1-HK2, and treated identically to the muscle cell lysate. Protein concentrations were determined using a Pierce BCA kit. Denaturation was performed by mixing the samples with buffer containing: 2.5% SDS and 50 μM β-mercaptoethanol and heated 30 min at 37 °C (Figs. 6m, 6n); or, with Sample Buffer containing 5% SDS and 100 mM dithiothreitol and heated 15 min at 65 °C (Figs. 6o, 6p). Protein markers were run on all gels). Gels were electrophoresed for 1.5 – 2h at 140 volts. Proteins were transferred to PDVF membranes overnight at 4 °C, 30 V. Membranes were stained with Ponceau S and washed, then incubated in TBST buffer with 2% nonfat dry milk for 1h at room temperature, followed by incubation with an SGLT2 primary antibody (ProteinTech 24654-1-AP ^53^, 1:500, 24 h; or Abcam ab37296 ^54^, 1:250, 48 h), washed 3×15 min in TBST buffer, incubated with secondary antibody (Licor IRDye @680 goat anti-Rabbit 925-68071, 1:10000)1.5 h at room temperature, washed in TBST buffer 3×15 min, and imaged (Odyssey Imager, LICORbio).

### Statistics and Reproducibility

GraphPad 10.2.3 and Sigma Plot 14 software were used for statistical analyses. Significant differences between means of normally distributed groups were evaluated by Paired Student’s T-Test. Mean values are given ± standard deviation. p values are reported as one-tailed with significance set at p < .05

## Data availability

The ICP-MS raw files are readable by proprietary software (MassHunter B.01.01 Build 123.10 Patch 3) and raw Excel files are available from the authors upon request. Other source data are provided with this paper. Individual uptake measurements from each muscle are included in the graphs.

## Acknowledgments

We thank Dr. Waldemar Priebe, Dept. of Experimental Therapeutics, Division of Cancer Medicine, University of Texas MD Anderson Cancer Center, for kindly providing custom synthesized ^13^C-substituted glucose derivatives. We thank Dr. Maria Czyzyk-Krzeska, University of Cincinnati, for providing the kidney cortical cell line. We acknowledge funding from the University of Cincinnati Physiology Research

Fund (JAH), University of Cincinnati Dean’s Innovation Research Fund (JH), NIH F31 DK132940 (NJN), and RO1 AR063710 (JAL).

## Author contributions

NJN, TLR, JAH, BM, and JAL designed measurements and analyzed results; NJN, LS, TZ, JH, and JAL measured results. PWC and MR designed and ran gene expression assays. IDF and WP synthesized ^13^C-αMDG. NJN, TLR, and JAH prepared the manuscript. All authors read and reviewed the final manuscript.

## Competing interests

The authors declare no competing interests.

## EXTENDED DATA

**Extended Fig. S1.**
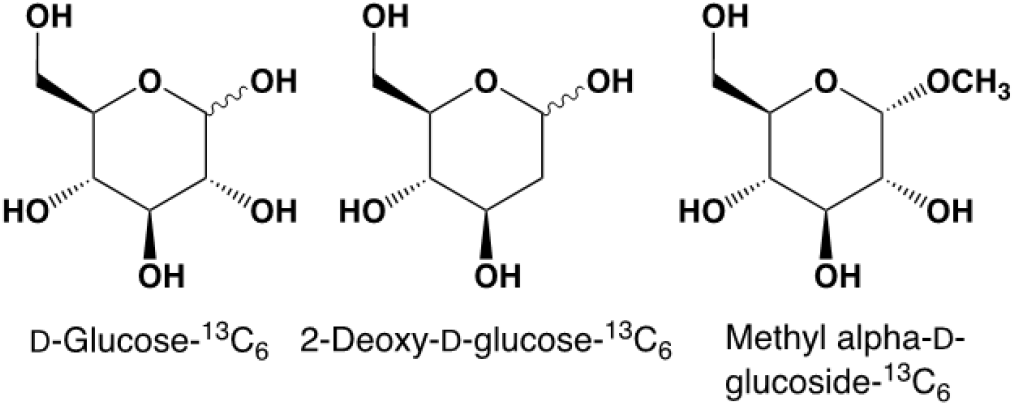
D-Glucose and its derivatives used in this study were isotopically labeled with ^13^C at all six carbon positions.

**Extended Fig. S2.**
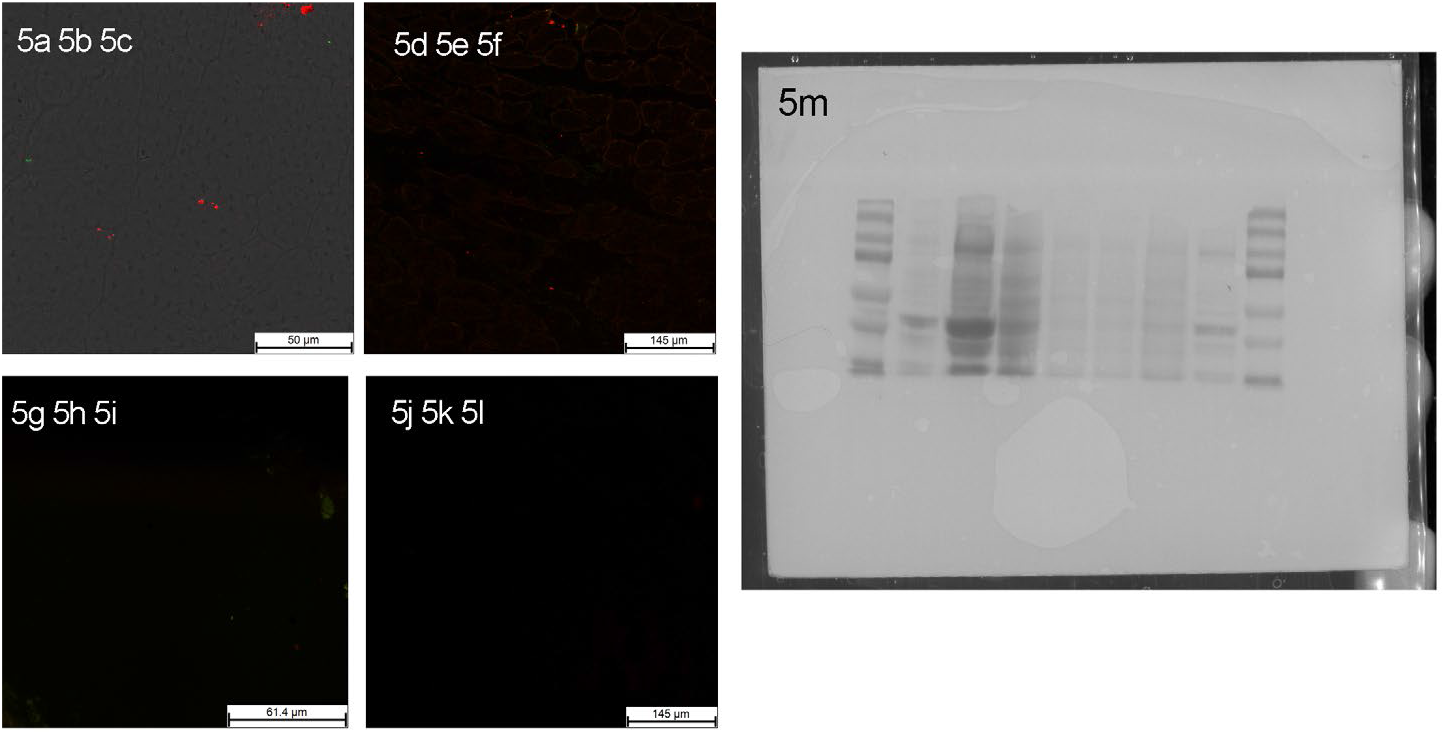
Control images for Fig 5a-5l, tissues incubated with secondary antibody only, no primary antibody. Ponceau S-stained membrane for Fig. 5m.

**Extended Table S1.**
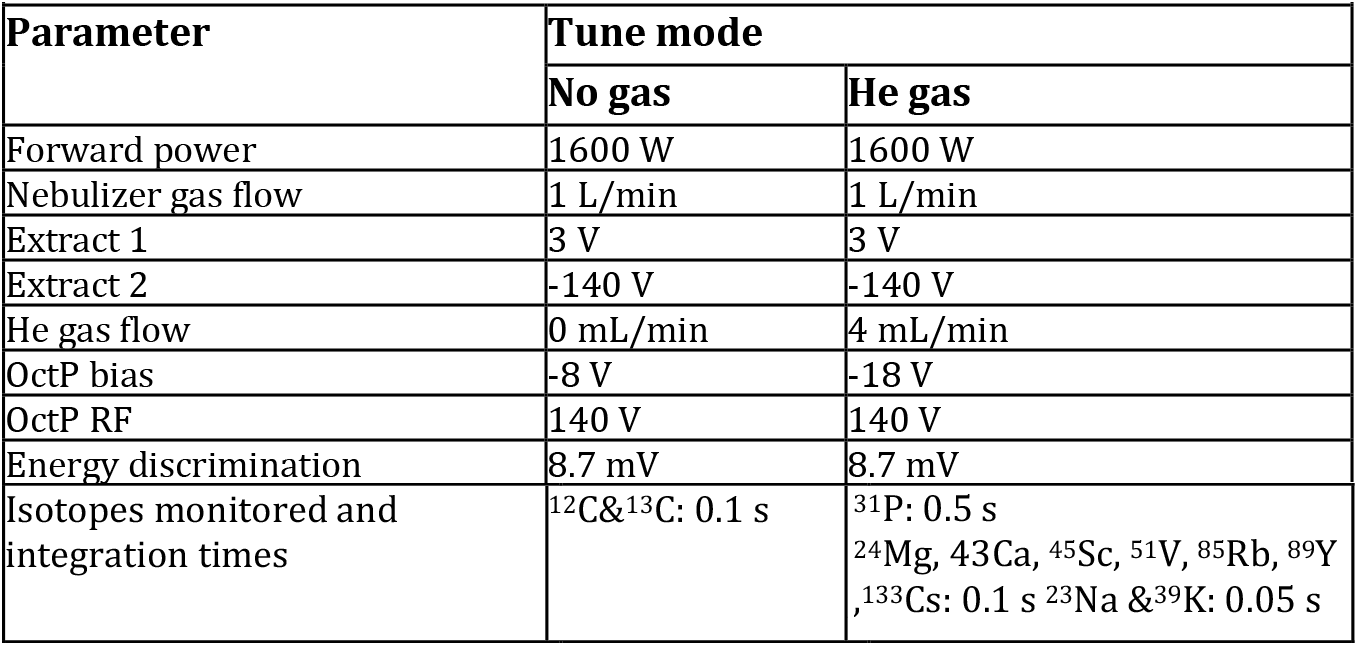
ICP-MS Tuning parameters.

| Parameter | Tune mode |  |
| --- | --- | --- |
|  | No gas | He gas |
| Forward power | 1600 W | 1600 W |
| Nebulizer gas flow | 1 L/min | 1 L/min |
| Extract 1 | 3 V | 3 V |
| Extract 2 | -140 V | -140 V |
| He gas flow | 0 mL/min | 4 mL/min |
| OctP bias | -8 V | -18 V |
| OctP RF | 140 V | 140 V |
| Energy discrimination | 8.7 mV | 8.7 mV |
| Isotopes monitored and integration times | $^{12}\text{C}$ & $^{13}\text{C}$: 0.1 s | $^{31}\text{P}$: 0.5 s<br>$^{24}\text{Mg}$, $^{43}\text{Ca}$, $^{45}\text{Sc}$, $^{51}\text{V}$, $^{85}\text{Rb}$, $^{89}\text{Y}$<br>$^{133}\text{Cs}$: 0.1 s $^{23}\text{Na}$ & $^{39}\text{K}$: 0.05 s |

**Table S2.** Numerical source data for Fig. 6e-6g. Rb and aMDG uptake in EDL muscles stimulated to contract or treated with salbutamol.

|  | Fig 6e |  | Fig 6f |  | Fig 6g |  |
| --- | --- | --- | --- | --- | --- | --- |
|  | **Rb** | **<sup>13</sup>C aMDG** | **Rb** | **<sup>13</sup>C aMDG** | **Rb** | **<sup>13</sup>C aMDG** |
|  | stim/stim<br>+phlorizin | stim/stim<br>+phlorizin | salbut/salbut<br>+empag | salbut/salbut<br>+empag | stim /stim<br>+empag | stim /stim<br>+empag |
|  | 1.029 | 0.85 | 0.51 | 1.30 | 0.97 | 0.84 |
|  | 1.791 | 0.706 | 0.54 | 1.56 | 1.06 | 0.67 |
|  | 0.876 | 0.63 | 0.50 | 2.13 | 0.88 | 0.71 |
|  | 0.922 | 1.157 |  |  | 1.14 | 0.79 |
|  |  | 0.939 |  |  | 0.95 |  |
|  |  |  |  |  | 1.03 |  |
|  |  |  |  |  | 1.07 |  |
|  |  |  |  |  | 1.05 |  |
|  |  |  |  |  | 0.84 |  |
| mean | **1.15** | **0.86** | **0.52** | **1.66** | **1.00** | **0.75** |
| SD | 0.43 | 0.21 | 0.02 | 0.42 | 0.10 | 0.07 |
| n | 4 | 5 | 3 | 3 | 9 | 4 |

## Notes

### Competing Interest Statement

The authors have declared no competing interest.

